# Boldine modulates glial transcription and functional recovery in a murine model of contusion spinal cord injury

**DOI:** 10.1101/2023.02.15.528337

**Authors:** Carlos A. Toro, Kaitlin Johnson, Jens Hansen, Mustafa M. Siddiq, Walter Vásquez, Wei Zhao, Zachary A. Graham, Juan C. Sáez, Ravi Iyengar, Christopher P. Cardozo

**Author notes:** **Correspondence: Carlos A. Toro**, /. **<u>Support</u>**, DOD SCIRP SC190031; VA RR&D 1I 50 RX002020; ANID1191329; VA RR&D 1IK2RX002781; NIH-GM 137056.

## Abstract

Membrane channels such as connexins (Cx), pannexins (Panx) and P2X_7_ receptors (P2X_7_R) are permeable to calcium ions and other small molecules such as ATP and glutamate. Release of ATP and glutamate through these channels is a key mechanism driving tissue response to traumas such as spinal cord injury (SCI). Boldine, an alkaloid isolated from the Chilean boldo tree, blocks both Cx hemichannels (HC) and Panx. To test if boldine could improve function after SCI, boldine or vehicle was administered to treat mice with a moderate severity contusion-induced SCI. Boldine led to greater spared white matter and increased locomotor function as determined by the Basso Mouse Scale and horizontal ladder rung walk tests. Boldine treatment reduced immunostaining for markers of activated microglia (Iba1) and astrocytic (GFAP) markers while increasing that for axon growth and neuroplasticity (GAP-43). Cell culture studies demonstrated that boldine blocked glial HC, specifically Cx26 and Cx30, in cultured astrocytes and blocked calcium entry through activated P2X_7_R. RT-qPCR studies showed that boldine treatment reduced expression of the chemokine Ccl2, cytokine IL-6 and microglial gene CD68, while increasing expression of the neurotransmission genes Snap25 and Grin2b, and Gap-43. Bulk RNA sequencing (of the spinal cord revealed that boldine modulated a large number of genes involved in neurotransmission in in spinal cord tissue just below the lesion epicenter at 14 days after SCI. Numbers of genes regulated by boldine was much lower at 28 days after injury. These results indicate that boldine treatment ameliorates injury and spares tissue to increase locomotor function.

## INTRODUCTION

Spinal cord injury (SCI) is a devastating form of neurotrauma that results in life-long disabilities. However, with the exception of physical rehabilitation, there is no clinically available cell-based or pharmacologic approach to improve sensory or motor function after SCI [1, 2]. The trauma results in a mechanical disruption of white and grey matter, shearing axons, crushing or shearing cell bodies, and causing damage or destruction of neural circuits [3]. The mechanical injury initiates a series of tissue responses that further damage surviving cells through increased levels of reactive oxygen species (ROS), excitotoxic effects of glutamate and activation of glia [3]. There then follows a cellular response that includes proliferation of astrocytes which surround the injury site, possibly to wall off the inflamed region [3]. Sprouting of surviving neurons occurs, which results ultimately in formation of relay circuits that support partial recovery of function [4, 5].

A growing body of evidence links membrane channels such as connexins (Cx), pannexins (Panx) and P2X_7_R channels to injury caused by tissue responses after SCI [6–10]. These channels share several biophysical properties including permeability to calcium ions. Cxs are a family of proteins that form hexameric pore structures called connexons, also referred to as Cx hemichannels (Cx HC), found largely in the cytoplasmic membrane but also in the membranes of organelles such as the mitochondria and endoplasmic reticulum. Individual Cxs proteins are named according to their molecular weights [11, 12] with connexin 43 (Cx43) is the most abundant Cx in the central nervous system where it is expressed in astrocytes, and to a lesser extent, in microglia [13]. Cx26 and Cx30 are also expressed on astrocytes of most species [13]. Cx HC are best known as the building blocks of gap junctions (GJ) that form when Cx HC on adjacent cells bind to one-another, forming a pore by which cells are electrically and chemically coupled. Cx HC also exist as cell surface pores where they connect cytosol and extracellular fluids. Open Cx HC permit NAD^+^, ATP, and glutamate to leave the cell, raising their extracellular concentrations [14, 15]. An inward flow of calcium through open Cx HC raises cytosolic calcium ion concentrations [11–13]. Cx HC are opened in response to various stimuli in physiological or pathological environments [16] such as pro-inflammatory cytokines and oxidative and metabolic stress [17–20]. While under physiological conditions, open Cx HC can modulate neuronal activity, in pathological situations they can also induce deterioration of cells and cell death [21, 22]. Pannexin 1 (Panx1) HC are permeable to ATP and are present in astrocytes and in neurons in which it is mostly localized to the post-synaptic zone. P2X_7_R allow calcium to enter the cell when bound by ATP, are localized to microglia and are present in lesser amounts on astrocytes and pre-synaptic neurons [23].

Previous studies of SCI in rodent models have demonstrated alterations in astrocytic Cx43 within the epicenter and penumbra of the lesion site [24, 25]. The transformation of astrocytes into reactive astrocytes increases Cx43 expression as a direct result of traumatic SCI [25]. Interestingly, Cx43 is the only Cx HC forming protein for which changes in expression are reported post-SCI [25, 26]. Several studies implicate Cx43 HC for a wave of ATP release into the extracellular space of tissues surrounding the lesion site as genetic ablation of Cx30 and Cx43 in astrocytes blunts ATP release and spares spinal cord tissue [7].

It should be noted that extracellular ATP binds to and activates P2X_7_R purinergic receptors on microglia resulting in maturation and release of IL-1ß [27] and that P2X_7_R blockers reduced inflammation and improved functional recovery after SCI [28]. In addition, binding of ATP to P2X_7_R channels adds to the inward calcium current and is linked to opening of Panx1 thereby further increasing release of ATP. Thus, P2X_7_R mediate a feed-forward amplification of signals originating as release of ATP through open Cx HC through coupling of P2X_7_R with Panx1 [29].

It was recently discovered that boldine, a naturally occurring alkaloid extracted from the leaves and bark of the Chilean boldo tree (*Peumus boldus*), blocks Cx43 and Panx1 HC without affecting GJ communication [14]. In a mouse model of Alzheimer’s disease, analysis of brain tissue showed that boldine prevented release of ATP and glutamate and lowered intracellular calcium concentrations [14]. These findings raised the question of whether boldine might reduce tissue injury and/or improve locomotor function after SCI. Here, we tested the hypothesis that administration of boldine beginning at 3 days after a moderate contusion SCI improves locomotor function and spare white matter.

## MATERIALS AND METHODS

### Animals

Use of live animals was conducted in accordance with PHS Policy on Humane Care and Use of Laboratory Animals and the Guide and was approved by the Institutional Animal Care and Use Committee at James J. Peters Veterans Affairs Medical Center (JJP VAMC) IACUC #CAR-20-11. Male and female C57Bl6 mice were purchased from Charles River and housed with controlled photoperiod (12/12 h light/dark cycle) and temperature (23–25°C) with ad libitum access to water and pelleted chow. Experiments using cultured astrocytes were performed following protocols approved by Ethics Committee of the Universidad de Valparaíso, Chile; ID: #CICUAL F-03.

### Experimental Design

A general summary of animal groups and sample sizes are provided in Table 1. At 4 months of age, male and female animals of similar weight (Supplemental Figure 1) were randomly assigned to SCI or a sham-SCI. Each group was then randomly assigned to boldine or vehicle-treated groups (Table 1). The experimental design and timeline for the study procedures is shown in Figure 1.

**Figure 1.**
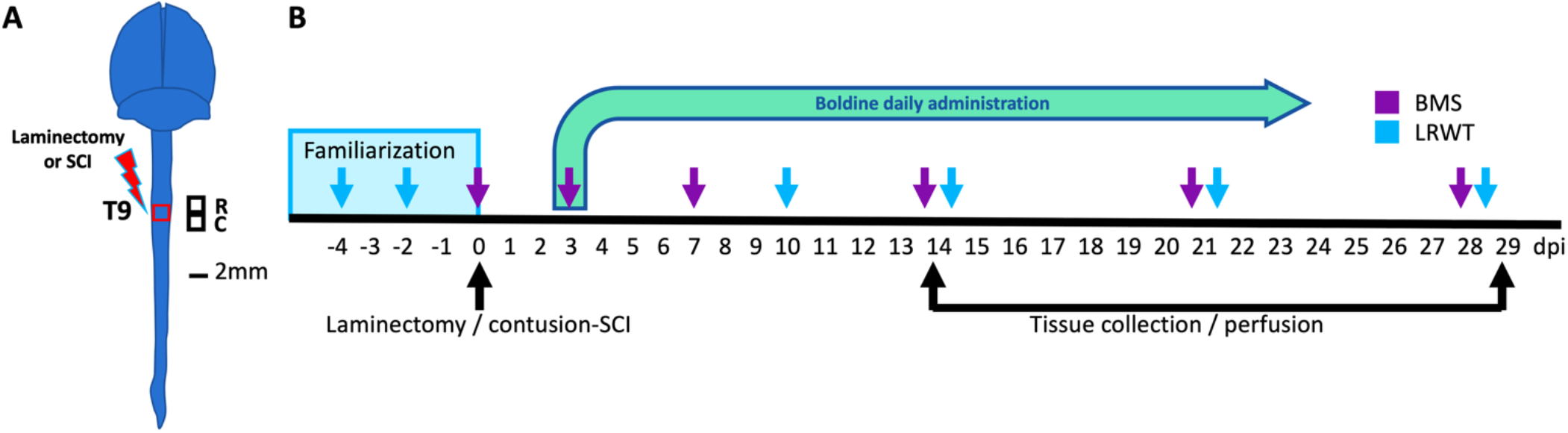
Experimental design. (A) Anatomical representation of the mouse brain and spinal cord showing the site of laminectomy or contusion SCI (red box) at the level of thoracic vertebrae 9 (T9). Black squares represent the locations of the rostral and caudal segments of spinal cord collected. These were used for histology, biochemistry and bulk RNA-seq studies. R: rostral; C: caudal. Scale bar, 2 mm. (B) Timeline for animal familiarization with handling, peanut butter and equipment, surgeries performed, boldine or vehicle administration, behavioral testing and timepoints for tissue collection and animal perfusion are depicted. BMS: Basso mouse scale; LRWT: ladder rung walk test.

**Table 1.** Experimental groups, procedures and treatments

| Table 1. Experimental groups, procedures and treatments |  |  |
| --- | --- | --- |
| Surgery/Treatment | Timepoints |  |
|  | 14-dpi | 28-dpi |
| Laminectomy (sham) / Vehicle | N= 8 Males; N= 6 Females | N= 8 Males; N= 6 Females |
| Laminectomy (sham) / Boldine | N= 8 Males; N= 6 Females | N= 8 Males; N= 6 Females |
| Contusion SCI / Vehicle | N= 10 Males; N= 8 Females | N= 10 Males; N= 8 Females |
| Contusion SCI / Boldine | N= 10 Males; N= 8 Females | N= 10 Males; N= 8 Females |

### Spinal Cord Injuries

A motor-incomplete contusion SCI was performed using an Infinite Horizons (IH) impactor (Precision Systems and Instrumentation) as previously reported [30]. Briefly, after induction of anesthesia by 3% isoflurane inhalation, animals were placed on heating pads warmed with recirculating water to 37 C. Hair was clipped over the cervical and thoracic spine areas. All animals underwent a laminectomy to expose the dura at level of thoracic vertebrae 9 (T9). The size of laminectomy was approximately 2 mm in diameter to allow the probe to impact the dura without touching any bone or other surrounding tissues. Randomized animals selected for the sham-SCI group had the vertebral muscle around the laminectomy site sutured to stabilize the spinal column, the incision site closed with 7 mm wound clips, and were returned to a clean cage on top of a warming pad. For animals selected for the SCI groups, they were placed on the clamping platform of the IH impactor under 3% isofluorane where their vertebral column was stabilized using the attached forceps to the IH clamping platform and received a 65 kdyne contusion SCI [31]. We chose this impact force as it results in a moderate-severe injury that allows for a partial recovery function over the timeframe of the study. Animals reach a maximal recovery and then plateau at around 28 days post injury. After the lesion is generated, the left-right symmetry was confirmed by detection of equal bruising on both sides of the dorsal median sulcus. Actual impact force and cord displacement values were recorded for each animal and group (Supplemental Figure 2). Muscle was closed using resorbable sutures and skin closed with 7-mm wound clips. Post-operative care took place in cages with Alpha-Dri bedding (Newco Distributors, INC.) over heating pads warmed with recirculating water for the first 72 h. All animals received pre-warmed lactated Ringers solution (LRS), carprofen and Baytril every 24 h for 5 d. Wound clips were removed at 10 days post injury (dpi). Urine was expressed manually twice a day by gentle pressure and massage of the bladder until spontaneous voiding. Animals were fully checked for signs of stress, urine scalds, and autophagia at least once a day for the length of the study.

### Boldine Administration

Boldine (Millipore Sigma) administration started at 3 dpi. It was prepared by dissolving boldine in a mix of DMSO and peanut oil (Sigma). This mixture was then added to peanut butter (PB) so that 1.0 g of total bolus could be given once per day at a dose of 50 mg/kg. The final concentration of DMSO was less than 2.5 %. Animals were familiarized with 1.0 gram of peanut butter for a week prior to surgery. All animals consumed 100% of their daily PB mix and continued to do so throughout the remainder of the study. Vehicle-treated SCI and laminectomy-only animals (shams) received daily equal amount of the PB mix without boldine.

### Behavioral Testing

Locomotor recovery after SCI was tested as previously reported [30] using the Basso Mouse Scale (BMS) open-field test [32], and the horizontal ladder rung walk test (LRWT) [33], at specified timepoints (Figure 1). BMS is a well-established method for evaluating the severity of impairments in locomotor function after SCI by scoring locomotor milestones using a 9-point scale, with 0 being completely hindlimb paralysis and 9 being a normal healthy gait. The LRWT is used to evaluate fine motor skills and coordinated stepping as mice attempt to cross a commercially available horizontal ladder and were evaluated as they attempted to cross [33]. For BMS, two blinded investigators independently scored the animals and the results were averaged for a final score. For LRWT, animals were recorded with a video camera located under the ladder that was moved manually to keep the animal in-frame and later reviewed by two blinded observers who recorded the number of correct steps and errors. Animals were familiarized with equipment used for each of these behavioral tests one-week prior to surgery. We performed all behavior tests during the animals’ dark phase of the light:dark cycle, which we set from 0600-1800 for this study, as they are normally more active in the dark.

### Tissue Harvest

At days 14 and 28 post SCI, 5 animals per group (Table 1) were randomly selected to undergo perfusion-fixation under a deep anesthesia following an intraperitoneal injection of ketamine (100 mg/kg) and xylazine (30 mg/kg) as previously reported [30]. Briefly, mice were euthanized by transcardial perfusion with sterile saline followed by injection if ice-cold 4% paraformaldehyde (PFA). After perfusion, spinal cords were removed and further post-fixed in 4% PFA for 72 h, transferred to a solution of 30% sucrose and stored at 4 C until cryostat sectioning. In addition, fresh spinal cord tissues from remaining animals of every group were collected after inducing deep anesthesia by inhalation of 3% isofluorane followed by decapitation. Spinal cord segments (~2 mm) containing half the lesion epicenter and the region immediately rostral or caudal, were collected and snap frozen in liquid nitrogen and then stored at −80 C for biochemical analysis; tissues from comparable areas of spinal cords of sham-controls were also collected.

### Detection of White Spared Matter

Transverse 10-micron sections of perfusion-fixed spinal cords were obtained with a cryostat (Leica) and used for immunofluorescence staining as previously reported [30]. Myelin was stained using Fluoromyelin according to the manufacturer’s protocol (FluoroMyelin green, ThermoFisher). To visualize spared myelin, sections of perfusion-fixed spinal cord were cut every 100 microns up beginning at 1 mm rostral to the injury epicenter and continuing through the epicenter, stopping 1 mm caudal to the epicenter. Sections were imaged with a confocal microscope and analyzed using ImageJ software (described below)

### Immunofluorescence Staining

Sections of perfusion-fixed spinal cord were used for immunofluorescence staining as previously reported [30]. Antibodies were used to detect the following proteins: GAP-43 (Abcam #ab16053) to label regenerating axons and identify sprouting; GFAP (Abcam #ab7260) to detect reactive astrocytes; and Iba1(Abcam #ab178846) for labeling macrophages and activated microglia. Secondary antibodies, Alexa Fluor 488-conjugated goat anti-mouse IgG (Abcam # ab150113) and Alexa Fluor 647-conjugated goat anti-rabbit IgG (Abcam # ab150079). Details in Table 2.

**Table 2.** Antibodies for immunohistochemistry

| Table 2. Antibodies for immunohistochemistry |  |  |  |  |
| --- | --- | --- | --- | --- |
| Antibody name | Isotype and host | Catalog number | Concentration | RRID |
| Anti-GAP43 antibody | IgG polyclonal, host rabbit | Abcam, ab16053 | 1:500 | AB_443303 |
| Anti-GFAP antibody | IgG polyclonal, host rabbit | Abcam, ab7260 | 1:500 | AB_305808 |
| Anti-Iba1 antibody | IgG Monoclonal, host mouse | Abcam, ab178846 | 1:500 | AB_2636859 |
| Alexa-488 anti-mouse | IgG polyclonal, host goat | Abcam, ab150113 | 1:2500 | AB_2576208 |
| Alexa-647 anti-rabbit | IgG polyclonal, host goat | Abcam, ab150079 | 1:2500 | AB_2722623 |

### Fluorescent in-vitro mRNA Hybridization

Fixed 10-micron spinal cord sections were mounted on Superfrost Plus slides (Thermo Fisher Scientific). Customized probes for GFAP, Cx43 and S100a8 were designed and provided by Advance Cell Diagnostics (Hayward, CA) for detection of mRNA. In situ hybridizations were performed according to the RNAscope Multiplex Fluorescent Reagent Kit v2 Assay protocol provided by the manufacturer and as described previously [34].

### Image Capture and Quantification

5 x 5 tiled images were obtained from stained sections using a 20X objective and a Zeiss 700 confocal microscope (Carl Zeiss). Blinded quantification was performed using ImageJ software (version 2.1.0/1.53c, National Institute of Health, USA) and integrated density of pixels was measured for each section and a mean value was calculated as previously described [30, 35]. Maximum background threshold was determined for each image and set for intensity quantification. Data are represented uding the mean ± SEM

### RNA Extraction, Reverse Transcription and qPCR

Total RNA was extracted from spinal cord segments that extended either ~2mm rostral or ~2mm caudal to lesion epicenter using TRIzol reagent (ThermoFisher) following the manufacturer’s instructions and methods previously described [30, 36]. Total RNA concentrations were determined by absorbance at 260 nm using a Nanodrop spectrophotometer (Thermo Scientific). RNA was reverse-transcribed into cDNA using Omniscript reverse transcriptase (Qiagen). PowerUp SYBR Green Master Mix (Thermofisher) was used to measure mRNAs of interest by qPCR. Primers were designed with help of the Primer Blast program from NCBI (Table 3). Formation of single SYBR Green-labeled PCR amplicons were verified by running melting curve analysis. Threshold cycles (CTs) for each PCR reaction were identified by using the QuantStudio 12K Flex software. To construct standard curves, serial dilutions were used from 1/2 to 1/512 of a pool of cDNAs generated by mixing equal amounts of cDNA from each sample. The CTs from each sample were referred to the relative standard curve to estimate the mRNA content per sample; the values obtained were then normalized for variations using peptidylprolyl isomerase A (Ppia) mRNA as the normalizing unit.

**Table 3.**
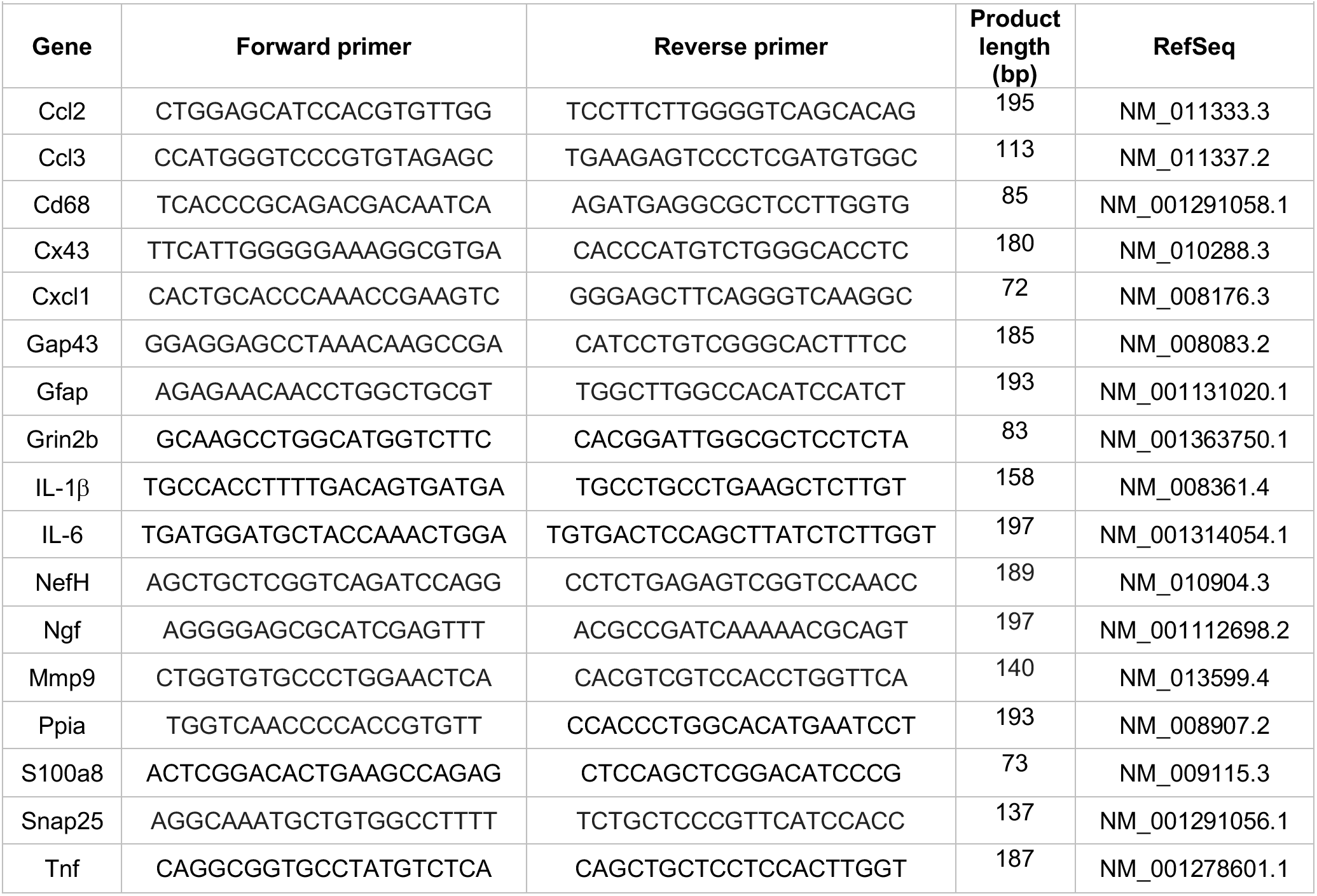
Primer sequence for RT-qPCR

### Transcriptomic Profiling by RNA Sequencing

We used total RNA extracted from spinal cord segments above and below the injury (~2 mm), from male mice at 14- and 28 days post SCI (N=3 per timepoint). RNA integrity was checked using the RNA 6000 Nano assay (Agilent). The sequencing library was prepared with a standard TruSeq RNA Sample Prep Kit v2 protocol (Illumina), as described previously [30, 37]. RNA libraries were sequenced on the Illumina HiSeq 2000 System with 100 nucleotide pair end reads, according to the standard manufacturer’s protocol (Illumina). For RNAseq data analysis, Star 2.5.4b and bowtie 2 2.1.0, samtools 0.1.7, Rsubread 2.10.5 and DESeq2 1.36.0 were used for read alignment to the mouse reference genome GRCm38.p6 using the ensemble gene annotation. At least 25 million reads were sequenced for each biological replicate (Supplemental Figure 6A). The percentage of reads that were successfully aligned to the mouse reference genome was between 84 and 89% (Supplemental Figure 6B). Differentially expressed genes (DEGs) were identified based on a maximum adjusted p-value of 10% (Supplemental Table 1). Up- and down-regulated gene sets were further interrogated using enrichR [38], with pathway enrichment analysis using Fisher’s Exact Test and the Gene Ontology Biological Processes 2018 library [39, 40] (Supplemental Table 2). Predicted pathways were ranked by significance. Significance p-values were transformed into -log10(p-values) and visualized as bar diagrams.

### Spinal Cord Astrocytes and HeLa cells to evaluate the activity of Cx26, Cx30 and P2X_7_R

Primary cultures of spinal cord astrocytes of three days old C57Bl6 mice were prepared using previously described methods [41]. Briefly, dissociated cells were resuspended in 10 ml DMEM with 10% FBS then transferred to a poly-lysine coated flask, place in an incubator at 37°C and 5% CO_2_, and allowed to adhere for 24 h. Flasks were then shaken at 200 rpm at 37 C overnight. The following day, non-adherent oligodendrocytes, neurons, and microglial were removed with aspiration and adherent cells were gently washed with DMEM containing 10% FBS, 10 mL of fresh DMEM with 10% FBS was added. Cells were allowed to recover and proliferate for 10 d before final seeding Hela cells transfected with mouse Cx30 or Cx26 were used (kind donation from Christian Giaume, College de France, Paris, France and Klaus Willecke, Life and Medical Sciences Institute, Molecular Genetics, University of Bonn, Bonn, Germany, respectively). The activity of Cx HCs was evaluated using the dye uptake method described previously [42, 43]. In brief, cells were plated onto glass coverslips and bathed with Locke’s saline solution (all concentrations in mM: 154 NaCl, 5.4 KCl, 2.3 CaCl_2_, 1.5 MgCl_2_, 5 HEPES, 5 glucose, and pH 7.4) containing 5 μM DAPI, a molecule that crosses the plasma membrane through large-pore channels, including Cx HC [24]. Since DAPI fluoresces upon its intercalation between DNA nucleotides, time-lapse recordings of fluorescent images were measured at regions of interest every 30 s for 13 min using a Nikon Eclipse Ti inverted microscope (Tokio, Japan) and NIS-Elements software. The basal fluorescence signal was recorded in cells only in Locke’s saline solution that contained divalent cations. Then, cells were exposed to Divalent Cation Free Solution (DCFS; Krebs buffer without CaCl_2_ and MgCl_2_), followed by 50 μM boldine and finally 200 μM La^3+^. Time-lapse fluorescence snapshot images were taken every 15 seconds. DAPI fluorescence was recorded in regions of interest using a Nikon Eclipse Ti microscope (Japan). Mouse P2X_7_R cDNA was cloned into pIRES-EGFP as previously described [44] (kindly donated by Dr. Claudio Acuña, Instituto de Química y Biología, University of Santiago) and transiently transfected into HeLa cells to evaluate the activity of P2X_7_R. Intracellular calcium was detected using Fura-2AM, a ratiometric dye. Cells were incubated in Krebs-Ringer solution (all concentrations in mM: 145 NaCl, 5 KCl, 3 CaCl_2_, 1 MgCl_2_, 5.6 glucose, 10 HEPES-Na, pH7.4) containing FURA2-AM dye (2 μM) for 45 min at room temperature. The calcium signal was then measured using a Nikon Eclipse Ti microscope equipped with epifluorescence illumination, and images were obtained by using a Clara camera (Andor) at 2 wavelengths of (λ) 340 nm and 380 nm, followed by calculating the ratio of fluorescence emission intensity after stimulation with each one of these two wavelengths. The activity of P2X_7_R was induced with 100 μM benzoyl ATP followed by the addition of 50 μM boldine. All above measurements were performed in ~20 cells per experiment in a total of at least four independent experiments.

### Statistical Analysis

Statistical evaluations were performed with one-way or two-way mixed model Analysis of Variance (ANOVA), as indicated on figure legends. Post-hoc comparisons were done using Tukey’s multiple comparison test. P values of less than 0.05 were considered significant. Statistical calculations were performed using Prism 9 software (Graphpad). Data are expressed as mean values are expressed as mean ± SEM.

## RESULTS

### Boldine administration promotes functional recovery after a motor-incomplete SCI

The effect of boldine on functional recovery after a thoracic motor-incomplete contusion SCI was evaluated by the open-field BMS test [32], and the horizontal LRWT [33] (Figure 2). Boldine or vehicle administration began at 3 dpi. All animals had maximum BMS scores before surgery (Figure 2A, B). As expected, shams treated with either vehicle or boldine presented maximum BMS scores at all timepoints (Figure 2A, B). SCI animals treated with boldine or vehicle presented similar BMS scores at day 3 (0.62 vs 0.61 for males; 0.14 vs 0.25 for females). However, scores for boldine-treated animals were significantly higher at 7 dpi in males when compared to SCI-vehicle mice by more than one-point on the BMS (3.41 vs 2.29). Boldine resulted in greater BMS mean scores in female mice compared to vehicle-treated animals at 7 dpi, though this did not reach our statistical threshold (2.25 vs 1.53). The differences between boldine- and vehicle-treated animals became greatest for both males and females at 14 dpi (6.56 vs 4.61 for males; 4.84 vs 3.53 for females) and stayed consistently elevated at 21 dpi (7.44 vs 5.57 for males; 5.89 vs 4.88 for females) and 28 dpi (7.94 vs 6.11 for males; 6.46 vs 5.00 for females) (Figure 2A, B). For LWRT, all sham animals made no step placement errors (Figure 2C, D), while the SCI-vehicle group demonstrated a numerically higher percentage of foot placement errors when compared to the SCI-boldine group at 10 dpi (46.4% vs 23.5% for males; 48.9% vs 34.8% for females). These differences met statistical thresholds for significance for both sexes at 14 dpi (37.4% vs 14.7% for males; 38.0% vs 21.6% for females), 21 dpi (30.2% vs 12.0% for males; 30.1 vs 16.0% for females) and 28 dpi (23.3% vs 8.9% for males; 22.0% vs 11.2% for females) (Figure 2C, D). Altogether, our behavioral data suggests that boldine administration starting 3 d after a 65 kdyne contusion SCI induces significant improvement of gross and fine locomotor function in both male and female mice.

**Figure 2.**
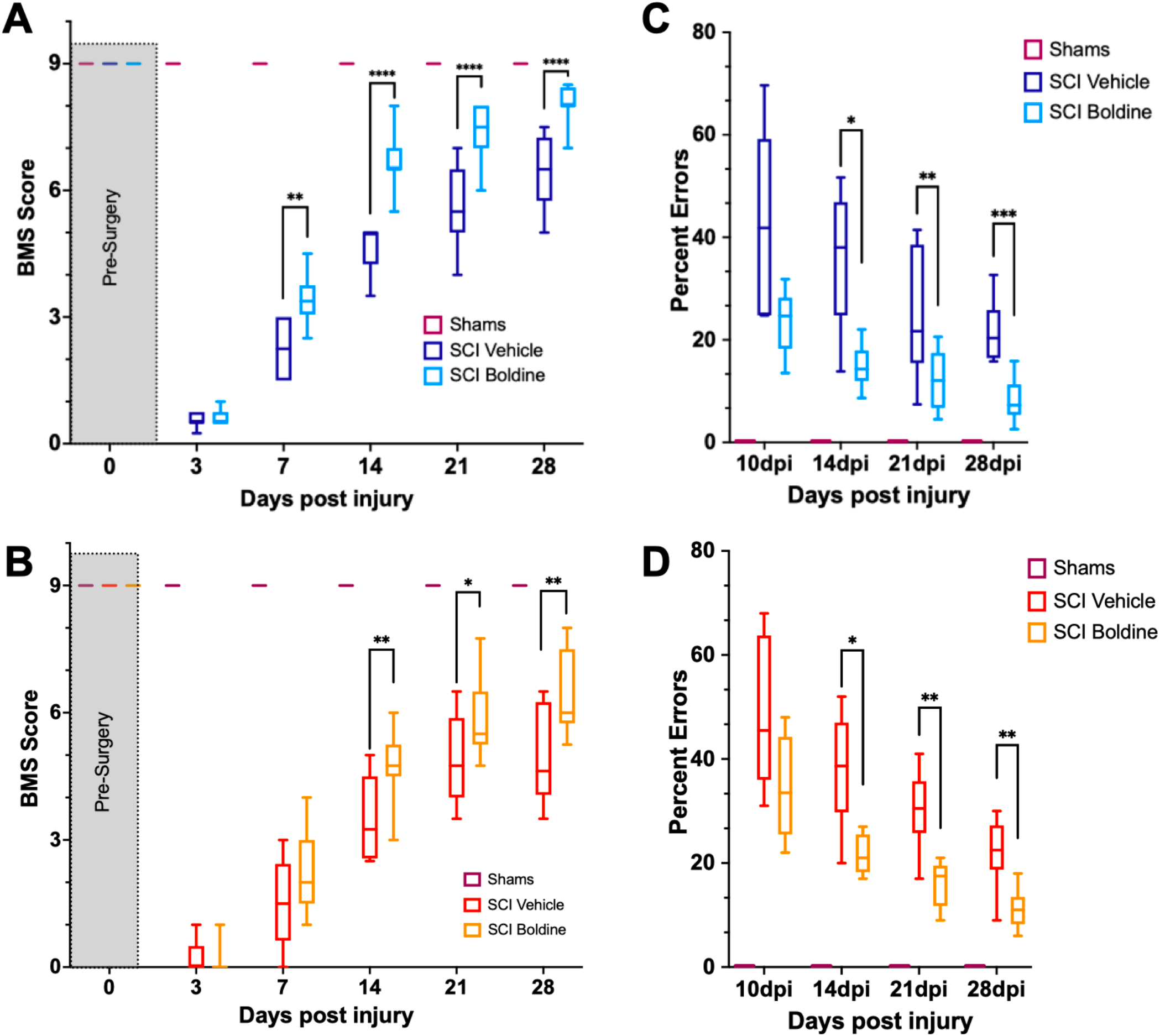
Boldine enhances functional recovery of C57BL/6 mice after contusion SCI. (A) BMS evaluation for shams, vehicle- and boldine treated male SCI mice. (B) BMS evaluation for shams, vehicle- and boldine treated female SCI mice. (C) LRWT scores for shams, vehicle- and boldine treated male mice. (D) LRWT scores for shams, vehicle- and boldine treated female mice. LRWT is expressed as percent of foot placement errors. Box-and-whisker diagrams represent the median, third quartile (upper edge) and first quartile (lower edge), and minimum and maximum values (whiskers) of the data. Statistical analysis was performed using a two-way mixed model ANOVA followed by a Bonferroni’s post hoc test. *p<0.05; **p<0.01; ***p<0.001; ****p<0.0001. N=10 males; N=8 females

### Boldine administration after moderate severity contusion SCI increases spared white matter and reduces lesion volume

To determine whether the overall size of the lesion was affected by boldine administration at 28 days post SCI, we performed fluorescent myelin detection (Fluoromyelin; ThemoScientific) and confocal imaging on spinal cord transverse sections from vehicle or boldine-treated animals every 100 μm including the injury epicenter the injury epicenter and locations rostral and caudal to it (Fig. 3A, B). Spared white matter was defined as the area stained with Fluoromyelin and the lesion size was measured as the disrupted area in each section. Area under the curve for spared white matter was significantly increased by boldine (Figure 3C) and the lesion volume was significantly reduced by boldine (Figure 3D) as compared to vehicle-treated animals. These data suggest that boldine reduces the lesion size and increase the abundance of white matter spared at 28 days after SCI.

**Figure 3.**
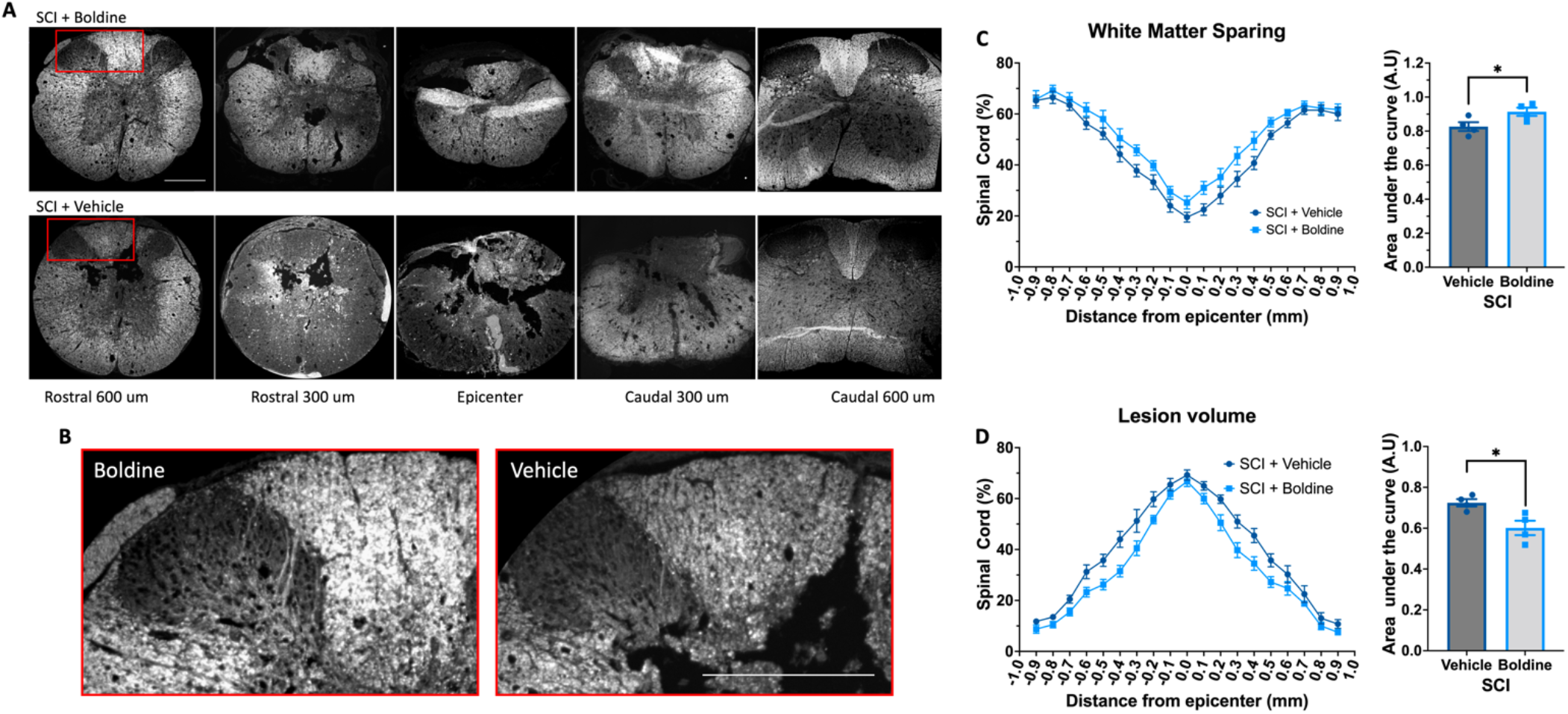
Boldine promotes sparing of white matter and reduces the lesion volume in male mice after SCI. (A) Perfusion-fixed spinal cords of boldine and vehicle treated SCI animals were cryo-sectioned at 28 days. Transverse sections were collected every 100 μm and stained with FluoroMyelin. Panel (A) shows representative images at the epicenter and at 300 and 600 μm rostral and caudal from epicenter for each group. (B) Higher magnification of regions within the red boxes in panel A depicting differences between boldine and vehicle treated groups at 600 μm rostral from the epicenter. Scale bar is 500 μm. (C-D) Transverse sections stained with FluoroMyelin were analyzed every 100 μm both, from 1 mm rostral to 1 mm caudal from the epicenter. (C) White matter sparing and (D) lesion site were detected and compared between boldine and vehicle groups. Area under the curve was calculated for each using ImageJ. Bar plots are presented as mean ± SEM. Statistical analysis was performed using unpaired t-test. *p<0.05. N=4 per group.

### Boldine reduces levels of reactive astrocytes and activated microglia and increases the expression of a protein involved in neuronal plasticity and growth cones after SCI

We further compared the injured spinal cord of vehicle or boldine treated animals at 14 dpi by immunofluorescence (IF) examination. This timepoint was chosen as the rate of functional recovery was most rapid post-SCI. To assess the effect of boldine on gliosis and after SCI, we probed sections taken just rostral to the lesion site for glial fibrillary acid protein (GFAP), a marker of reactive astrocytes [45] and Iba1, a marker associated with macrophage and microglia activation [46, 47]. Fluorescent signals of both GFAP and ionized calcium binding adaptor molecule 1 Iba-1 were significantly reduced by boldine (Figure 4A, B, respectively). Moreover, to investigate changes in expression of proteins related to neuronal plasticity and axon growth, we performed immunostaining of growth-associated protein 43 (GAP43) [48]. We show GAP43 fluorescence was significantly higher in sections rostral to the injury site of boldine-treated mice compared to vehicle-treated animals (Figure 4C). These data suggest that boldine reduces glial reactivity while stimulating plasticity of spared fibers 14 dpi after SCI.

**Figure 4.**
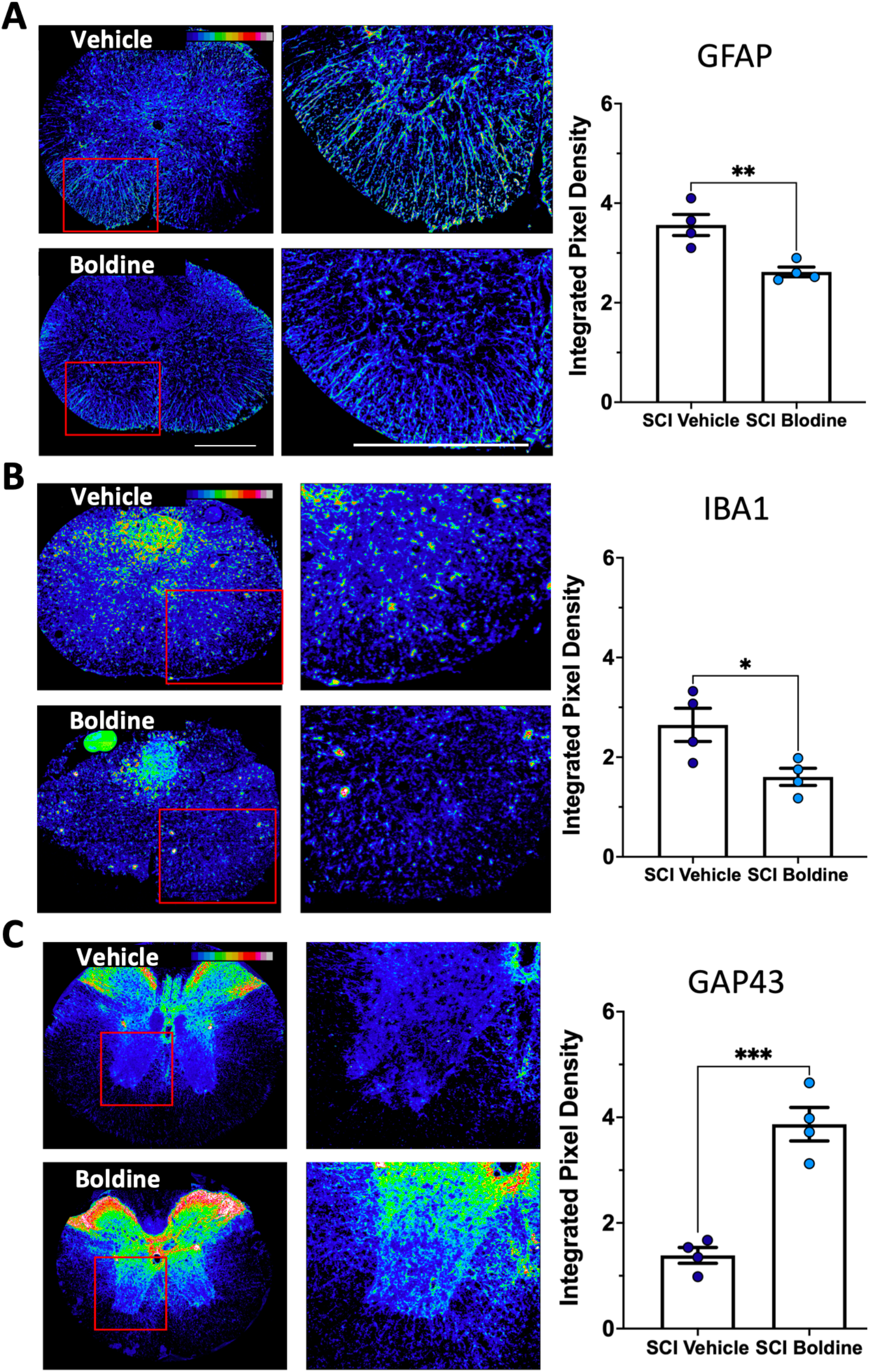
Boldine alters levels of protein markers for reactive astrocytes, activated microglia and neurite outgrowth after SCI. 10 μm transverse spinal cord sections surrounding the lesion site collected at 14 dpi were immunostained and analyzed to detect changes between boldine and vehicle treated groups using: (A) reactive astrocyte marker GFAP; (B) activated macrophage / microglia marker Iba1; (C) axonal growth cone marker GAP43. Immunofluorescence intensity is depicted using a 16-colorized scale from black/blue (lowest) to red/white (highest) as shown in the top right corner of the first figure for each panel. Blinded quantification of immunolabeling, at the right of each panel, was performed by evaluating integrated pixel density using ImageJ and comparing the results between boldine and vehicle groups. Images displayed are representative examples from N of 4 SCI male mice, per group. Objective 20X. Scale bar, 500 microns. Bar plots show means ± SEM. Statistical analysis was done using unpaired t-test. ***p<0.001, **p<0.005, *p<0.01.

### Boldine blocks Cx HC-mediated dye uptake in cultured embryonic spinal cord astrocytes

It is known that boldine blocks Cx43 and Panx1 HC in the brain [14]. To demonstrate that boldine blocks HC present on the surface of spinal cord astrocytes, primary cultures of spinal cord astrocytes were maintained in divalent-cation-free culture media in the presence of DAPI, which is taken up by live cells through HC [14], with or without 50 μM of boldine. Boldine significantly reduced uptake of DAPI during time-lapse imaging (Supplemental Figure 3) indicating that it blocked open HC in these cells.

### Boldine blocks Cx26, Cx30 and P2X_7_R channels

Multiple Cx other than Cx43 are expressed in the central nervous system, including Cx26 and Cx30 [49, 50]. Therefore, we tested if boldine blocks dye uptake in open Cx26 and Cx30 HC using transfected HeLa cells. Dye uptake studies demonstrated that boldine slowed the rate of dye uptake through both Cx26 and Cx30 (Supplemental Figure 4). Addition of lanthanum, a well-stablished Cx HC blocker [51], did not further reduce the rate of dye uptake, confirming that boldine acted by blocking Cx26 and Cx30 HC. Because the effects of boldine on P2X_7_R uptake has not been tested, additional cytoplasmic calcium signal studies were performed. The data showed that 50 μM boldine also blocks P2X_7_R (Supplemental Figure 5).

### Boldine modulates spinal cord transcriptomic profiles after SCI

An unbiased understanding of the cellular and molecular mechanisms responsible for the differences observed in locomotor recovery between SCI animals treated with vehicle versus boldine was explored through bulk RNA-seq followed by a bioinformatic analysis. Total RNA was extracted from spinal cord segments rostral and caudal from the lesion epicenter (~4 mm) at 14 dpi, time when functional recovery differences were maximal in between treatments, and at 28 dpi, when functional recovery had reached a plateau in maximal locomotor function. Differentially expressed genes (DEG) were identified and the magnitude of change and number of DEGs that differed between boldine and vehicle treated SCI animals was reported (Figure 5, Supplemental Figure 7). The data showed unique differences within the spinal cord segment at 14 dpi: while we found only 6 and 18 up- and downregulated genes above the lesion in boldine-treated animals (Figure 5A), we identified 426 and 913 up- and downregulated genes (Figure 5B) below the lesion in boldine treated SCI mice, respectively.

**Figure 5.**
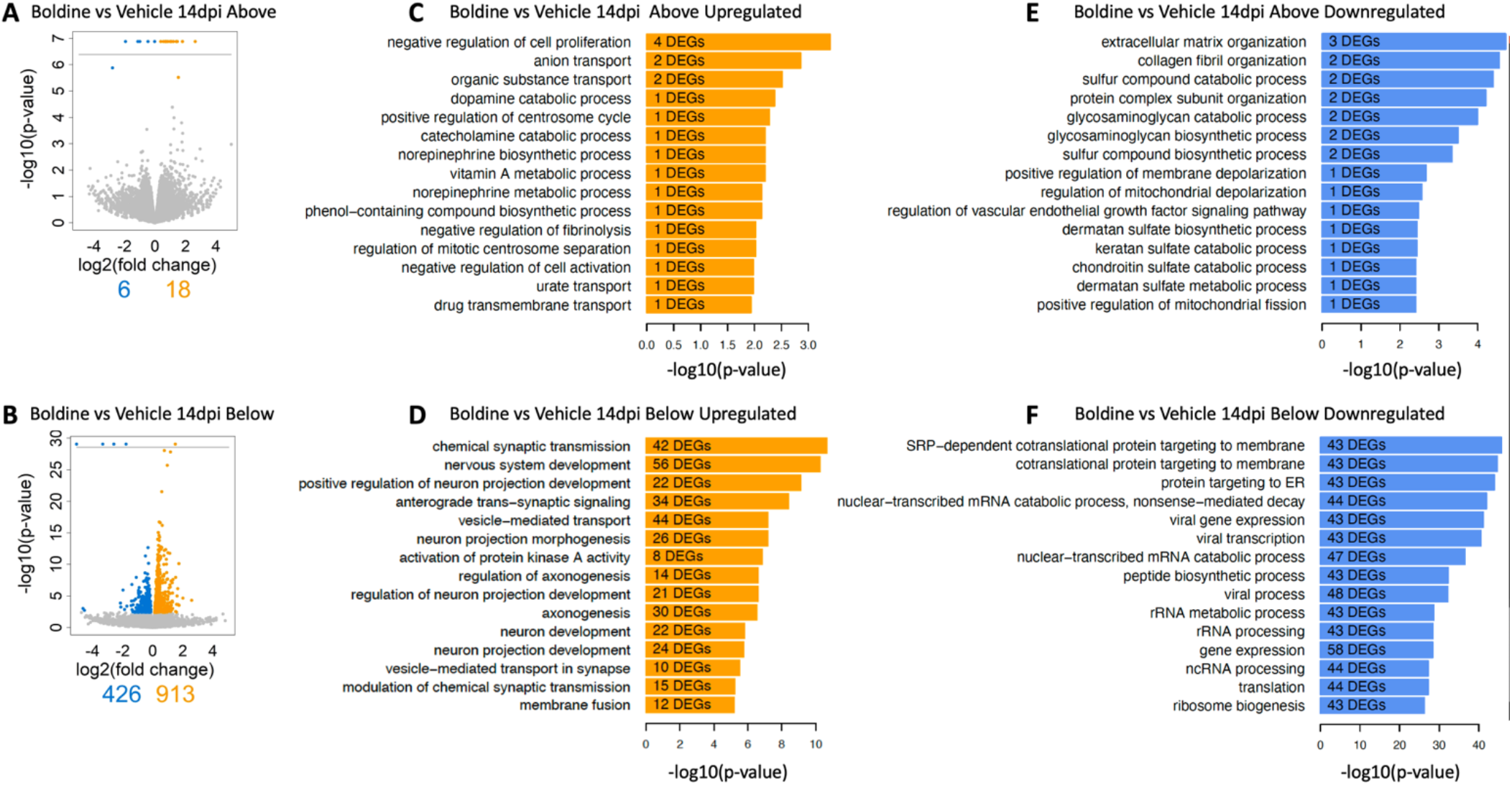
Boldine alters the injured spinal cord transcriptomic profile mostly below the lesion at 14 days after injury. Segments of spinal cord tissue collected at 14 dpi from mice with contusion SCI treated with either boldine or vehicle were subjected to bulk-RNA sequencing. Drug-induced DEGs (FDR 10%) were identified between boldine and vehicle treated SCI animals either above (A-C) or below (D-F) the injury at 14 days post injury. (A and D) Blue and orange dots indicate significantly up- or downregulated genes, respectively. DEGs predicted with a p-value of 0 and consequently -log10(p-value) of infinity are visualized above the horizontal line. (B, C, E and F). Up-regulated and downregulated genes for a particular condition were subjected to pathway enrichment analysis using Gene Ontology Biological Process and Fisher’s Exact test, followed by ranking of the predicted pathways by significance. The top 15 ranked pathways are shown for all lists that describe differences between treatments above or below the lesion site: (B) Boldine (vs vehicle) above the lesion upregulated, (C) Boldine (vs vehicle) above the lesion downregulated, (E) Boldine (vs vehicle) below the lesion upregulated and (F) Boldine (vs vehicle) below the lesion downregulated. Numbers of DEGs observed in each particular pathway are shown within the bar for that pathway.

To understand more in depth the biology associated with changes in numbers of DEGs, we identified biological processes represented by up- and downregulated genes above and below the lesion site (Figure 5C-F). While the top 15 predicted pathways above the lesion did not contain any pathways related to recovery of synaptic function or neuronal repair for any list of DEGs, the top upregulated pathways below the lesion for boldine treated animals at 14 dpi focused on functions related to neuronal development, synaptic transmission and axonogenesis (Figure 5D). In contrast, the pathways that were downregulated below the lesion included functions such as catabolic and biosynthetic processes (Figure 5F).

### Expression of pro-inflammatory molecules and genes involved in gliosis and neuronal function and plasticity are regulated by boldine after SCI

To further investigate the effect of boldine in changes of mRNA levels after SCI, differences in targeted gene expression between vehicle and boldine treated animals were evaluated using spinal cord segments (~2 mm) right above and below the lesion site at 14 dpi. RT-qPCR experimentation revealed that levels of pro-inflammatory molecules Ccl2 (Figure 6A), IL-6 (Figure 6B) and S100a (Figure 6C) were significantly greater in SCI animals treated with vehicle. Interestingly, these changes were not detected in boldine samples (Figure 6). Moreover, there was a similar trend observed for genes that express Ccl3, Tnf, Cxcl1 and IL-1b (Supplemental Figure 8 A-D, respectively), all cytokines and chemokines involved in inflammatory responses [52].

**Figure 6.**
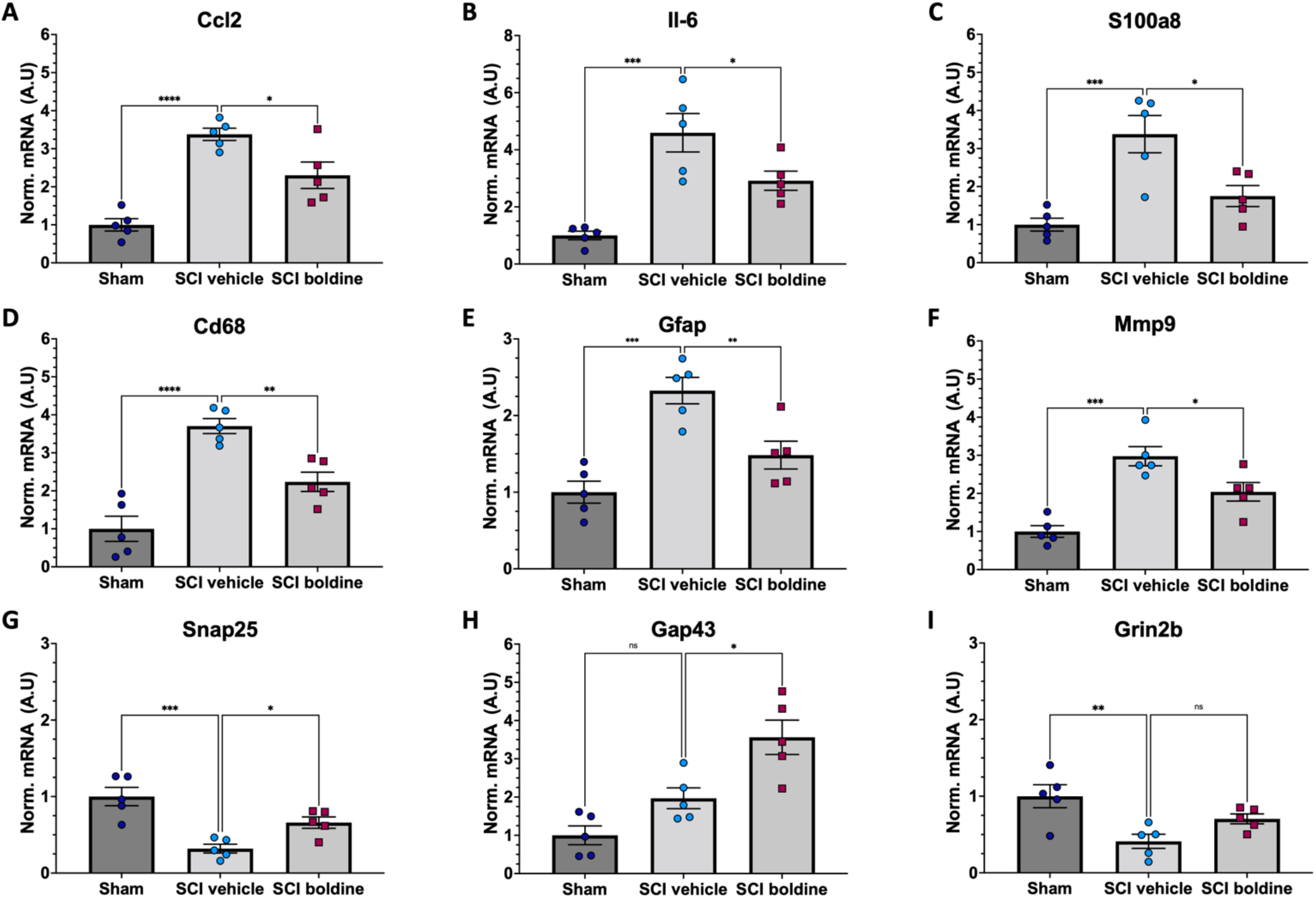
Effect of boldine in the modulation of mRNA levels of pro-inflammatory genes and genes related to gliosis, and neuronal plasticity and synaptic function after SCI. RT-qPCR was performed using total RNA isolated from 4 mm spinal cord segments spanning T7 to T11 (Figure 1) from sham and SCI mice at 14 dpi. Levels of (A) Ccl2, (B) Il-6, (C) S100a8, (D) Cd68, (E) Gfap, (F) Mmp9, (G) Snap25, (H) Gap43, and (I) Grin2b were detected for laminectomy-only (Sham), and for SCI animals treated with vehicle (SCI vehicle) or boldine (SCI boldine). Data are expressed as arbitrary units (A.U.) after normalizing to SCI vehicle samples. Bar plots show mean ± SEM. Statistical analysis was performed by one-way ANOVA followed by Tukey’s multiple comparisons test *p<0.05; **p<0.005; ***p<0.001; ****p<0.0001. N=5

We then evaluated changes in the expression of Cd68, a maker of activated microglia [53], Gfap, a highly expressed in reactive astrocytes [45], and Mmp9 which encodes for an enzyme with degradation of extracellular matrix activity and protection of motor neuron death [54]. Levels of Cd68, Gfap and Mmp9 were significantly increased in samples from injured spinal cords around the lesion. However, these changes were not observed in samples from animals treated with boldine (Figure 6 D-F, respectively). Levels of Cx43 HC also increased after injury, but were not significantly decreased in samples from boldine treated animals (Supplemental Figure 8E).

We also tested for genes that encode for SNAP25, involved in synaptogenesis [55]; Gap43 involved in axonal growth and plasticity [48], and for the NMDA receptor subunit, Grin2b [56]. Interestingly, levels of Snap25 and Gap43 are significantly higher in samples from animals treated with boldine as compared to vehicle treated samples, and a similar trend is observed for Grin2b although not significant (Figure 6 G-I). In addition, we evaluated changes for genes NefH and Ngf, also involved in axonal function, growth, maintenance and neuron survival [57–60] although changes between vehicle and boldine samples were not significant and only showed a trend (Supplemental Figure 8F,G).

We further performed RNAscope *in-situ* hybridization assay to detect mRNA transcripts and validate the changes observed for Gfap, S100a8, and Cx43 using transverse cryo-sections of spinal cords immediately below the lesion epicenter of boldine and vehicle treated SCI animals at 14 dpi. Our results showed, as expected, significantly higher levels of GFAP and S100a8 in vehicle compared to boldine-treated SCI animals (Figure 7 A-E). However, levels of Cx43 transcripts remain similar at this timepoint (Figure 7 D).

**Figure 7.**
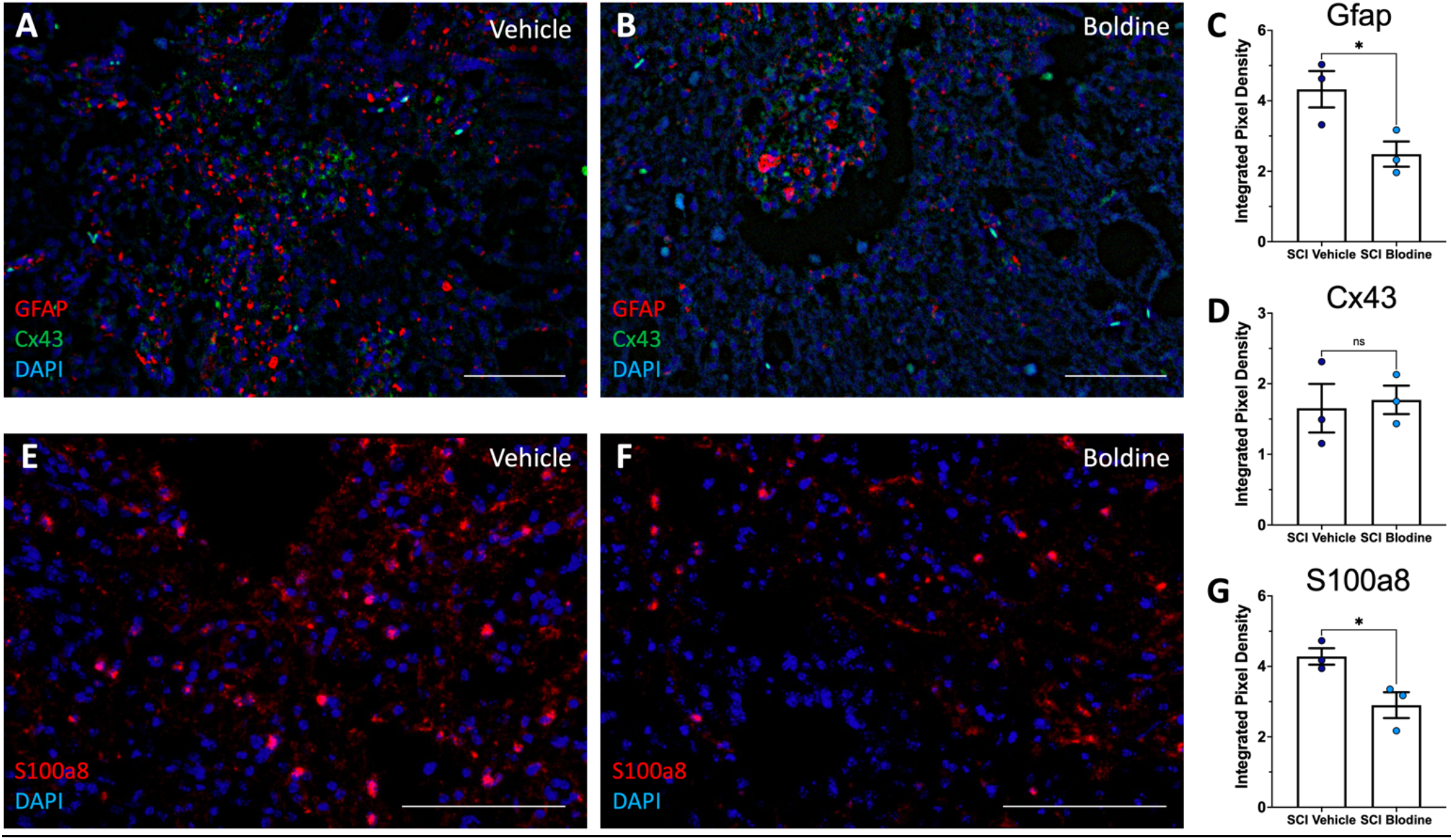
Boldine modulates mRNA levels of Gfap and S100a8. Transverse cryosections of spinal cords collected at 14 dpi were processed for visualization of mRNA transcripts for Gfap (A, B; red), Cx43 (A, B; green) and S100a8 (E, F; red) and using RNAScope and then visualized by confocal microscopy. Representative images show labeling of Gfap and Cx43 and S100a8 from vehicle treated animals (A, E) and boldine treated animals (B, F). Staining intensity was quantified and plotted as integrated pixel density for Gfap (C), Cx43(D) and S100a8 (G). Scale bar, 500 microns. Plots show mean value ± SEM. *, p < 0.05, unpaired 2-tailed t-test. N=3.

## DISCUSSION

The above experiments tested if oral administration of boldine improved locomotor function after a moderate contusion SCI in mice. The rationale for testing boldine stemmed from early work showing the critical role of Cx43 HC in the wave of ATP release that follows contusion SCI and which drives secondary injury through binding to and activation of P2X_7_R channels [7, 61] and from findings that boldine blocks open Cx43 HC [14]. The major conclusion of this study is that boldine resulted in increased locomotor function compared to vehicle-treated SCI animals as assessed by the BMS and LRWT. BMS test results were improved by as much as 2 points on this 9-point scale. Numbers of errors during LRWT were reduced by up to one half by boldine suggesting substantial improvements in fine motor skills. Additional literature supports a feedforward pathway that amplifies the effects of release of ATP through open Cx HC wherein binding of ATP to P2X_7_R opens Panx1 HC further increasing ATP release and inward calcium currents. Thus, the ability of boldine to block P2X_7_R and Cx26, Cx30 suggest that boldine blocks multiple channels involved in this feed-forward inflammatory pathway.

The experimental design incorporated a delayed start of boldine administration. One reason for this was that several prior studies established that blocking CxHC using the mimetic peptide 5 or inhibitory monoclonal antibodies improved locomotor function when treatment was begun soon after the contusion [62–64]. It was not, however, known if substances that block CxHC could improve function when begun at later times after injury. The data reported here provide evidence that in mice with contusion SCI, boldine improved function even when its administration is delayed for 3 dpi. An unanswered question is whether boldine would further improve locomotor function after SCI if started at 1 h or 1 d after contusion. Studies showing that monoclonal antibodies administered intrathecally at 1 month after contusion did not improve locomotor function suggest that there is a time beyond which Cx HC blockers do not change the course of SCI [64]. Perhaps by this time, either all Cx HC have closed or the cells expressing them are no longer situated near neurons and axons such that they are unable to affect health of these cells. More experimentation is needed to map out the temporal and spatial localization of open Cx HC after SCI.

Spared white matter is a well-established determinant of locomotor function after contusion SCI [65]. Findings that boldine treatment resulted in greater Fluoromyelin staining support white matter sparing as one mechanism by which boldine increased locomotor function. Interpretation of the effect of boldine on white matter should include some caveats. Specifically, BMS scores were indistinguishable at 3 dpi when comparing boldine-treated and vehicle-treated SCI groups but were higher in boldine-SCI mice compared to vehicle-SCI mice 4 days later. This suggests that initial damage to white matter was most likely comparable between boldine and vehicle groups and that boldine reduced subsequent white matter damage incurred beyond 3 days after the contusion. While it is well-accepted that demyelination and axon dieback continue for days to weeks after SCI [66], effects of treatments or genetic modifications on loss of white matter beyond the first few minutes to hours after the SCI is a relatively understudied area of research. Our data provide experimental support for the possibility that glia responses to SCI could continue to stress axons days after injury and that the mechanisms can be inhibited by boldine.

The mechanisms for protection by boldine against delayed axon loss cannot be determined with certainty from our data. It is notable that immunostaining for the reactive astrocyte marker GFAP and the microglial marker Iga1 was increased in white matter in vehicle-SCI compared to boldine-SCI mice. It is quite likely that these cell types remain activated after SCI for days, contributing to release of reactive oxygen species, excitotoxin such as glutamate and ATP, and pro-inflammatory cytokines. This interpretation is consistent with reduced activation of these cell types as assessed by immunofluorescence staining for protein markers, possibly lowering stresses on nearby axons. While logical, this interpretation requires experimental validation.

Rewiring of the remaining neural circuitry is critical to functional recovery after SCI. The current understanding of such rewiring is that a key step requires formation of axon branches arising from axons above the site of axon injury; these branches project to regions in the brain or spinal cord where they synapse with cell bodies of neurons that project axons through spared white matter to cell bodies below the lesion [67–69]. Below the lesion, additional axon branches may be formed to synapse with alpha motor neurons either directly or through interneurons [69]. Some of the relay circuits formed in this way have been experimentally shown be responsible for recovery of locomotor function [69]. Formation of these relay circuits must require specific gene expression programs to support axon growth and remodeling of extracellular matrix. Our data do not permit us to determine whether boldine increased formation of relay circuits. However, the data do provide indirect evidence that boldine increased remodeling of local neural circuitry. Specifically, spinal cords of boldine-SCI mice demonstrated greater immunolabeling for protein markers of axon growth cones, and higher levels of mRNA for synaptic function. Further studies using tract tracing and synaptic silencing are needed to understand whether boldine indeed alters formation of relay circuits.

A surprising finding was that bulk RNA sequencing showed few DEGs in tissue samples just above the lesion but many DEG below it; DEG below the lesion represented multiple gene ontologies involved in neurotransmission. One interpretation of these findings is that because axons are tiny relative to cell bodies, the changes in mRNA levels responsible for increased axonal expression of GAP-43 are not detected against the overall levels and magnitude of changes of mRNAs from whole tissue homogenate. In contrast, below the lesion, there may be many neurons responding to new synaptic inputs received from new axon branches eliciting large changes in gene expression programs in these cell bodies. This interpretation is consistent with findings from recent studies from the Levine lab have shown that extensive changes in expression of genes occur in selected populations of neurons in lumbar spinal cord after a severe thoracic contusion SCI [70]; Further study is needed to define the impact of boldine on neurons below the lesion after SCI.

In conclusion, the findings of this study show that boldine spares white matter and improves locomotor function in mice with a moderate severity mid-thoracic spinal cord contusion. Limitations include an incomplete understanding of molecular mechanism by which boldine achieves these effects and uncertainty regarding effects of time after SCI at which boldine is administered on its ability to improve locomotor function after SCI. It will be exciting to learn results of experiments addressing these limitations.

## Supporting information

Supplemental RNAseq data1

Supplemental RNAseq data2

## AUTHOR CONTRIBUTIONS

Conceptualization, CPC, JCS and CAT; Methodology, CAT, KJ, MS, WV, JCS and JH; Software, JH, RI; Investigation, CAT, WZ, GZ, MS, WV and CPC; Formal Analysis, CAT, JH, KJ, WV, MS; Resources, CPC, JCS and RI; Writing – Original Draft, CAT and CPC; Writing – Review & Editing, CAT, WZ, MS, JH, ZG, JCS, RI, JCS and CPC; Visualization, CAT, MS, JH and JCS; Funding Acquisition, CPC, and RI; Supervision, CPC, JCS and RI.

## ACKNOWLEDGEMENTS

To Drs. Miguel Gama-Sosa and Rita DeGasperi for helpful discussions related to technical aspects of the studies reported.

## DATA AVAILABILITY STATEMENT

The data are available upon written, reasonable request. Gene Expression Omnibus (GEO) data accession GSE220907.

