## Supplemental Figures for "Boldine modulates glial transcription and functional recovery in a murine model of contusion spinal cord injury"

SUPPLEMENTAL MATERIAL

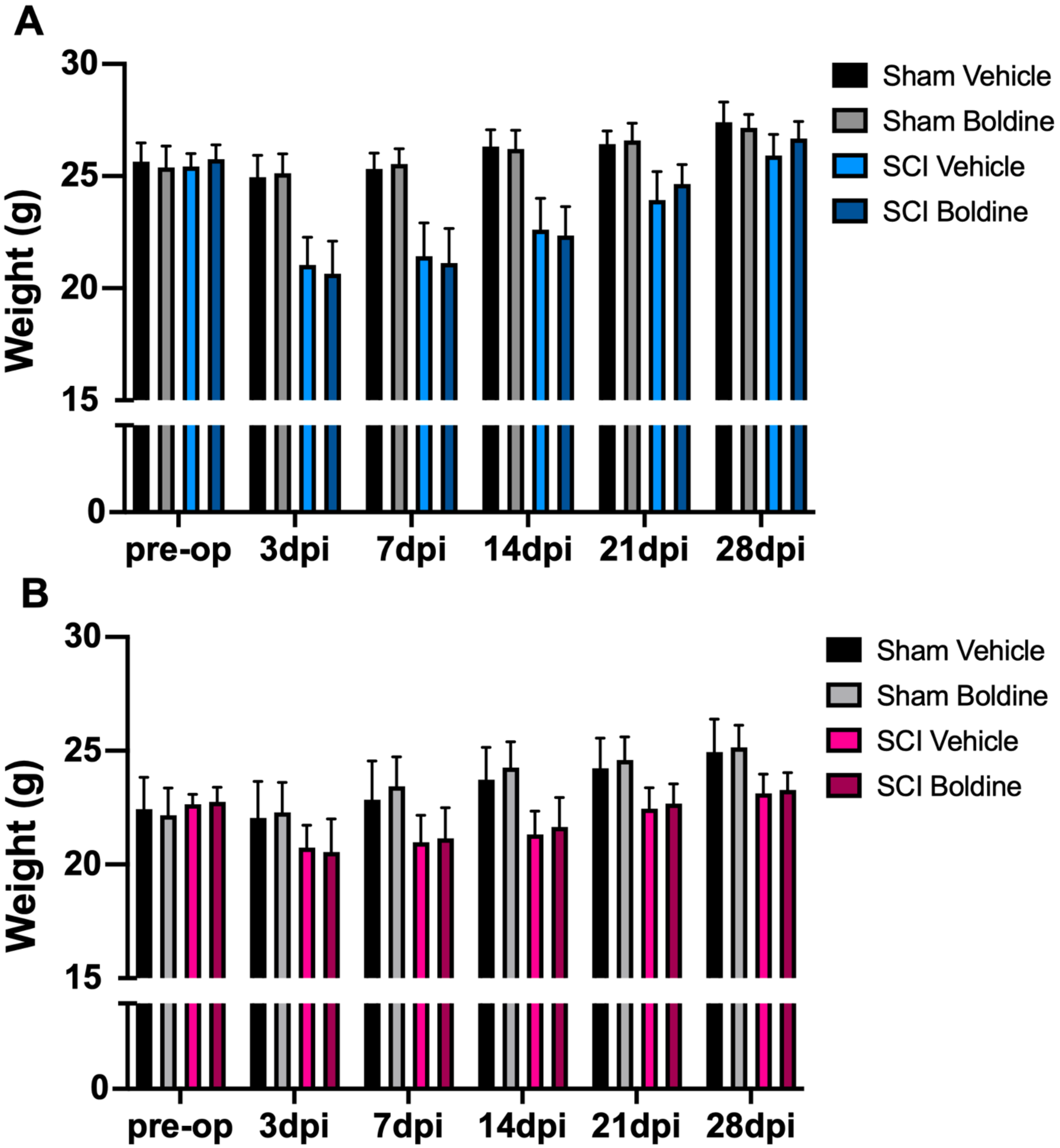


Supplemental Figure 1. Body weights were determined on the day of surgery prior to making an incision (pre-op) and at the specified times shown on the X-axis for male (A) and female (B) mice. Data are shown as mean values ± SEM.

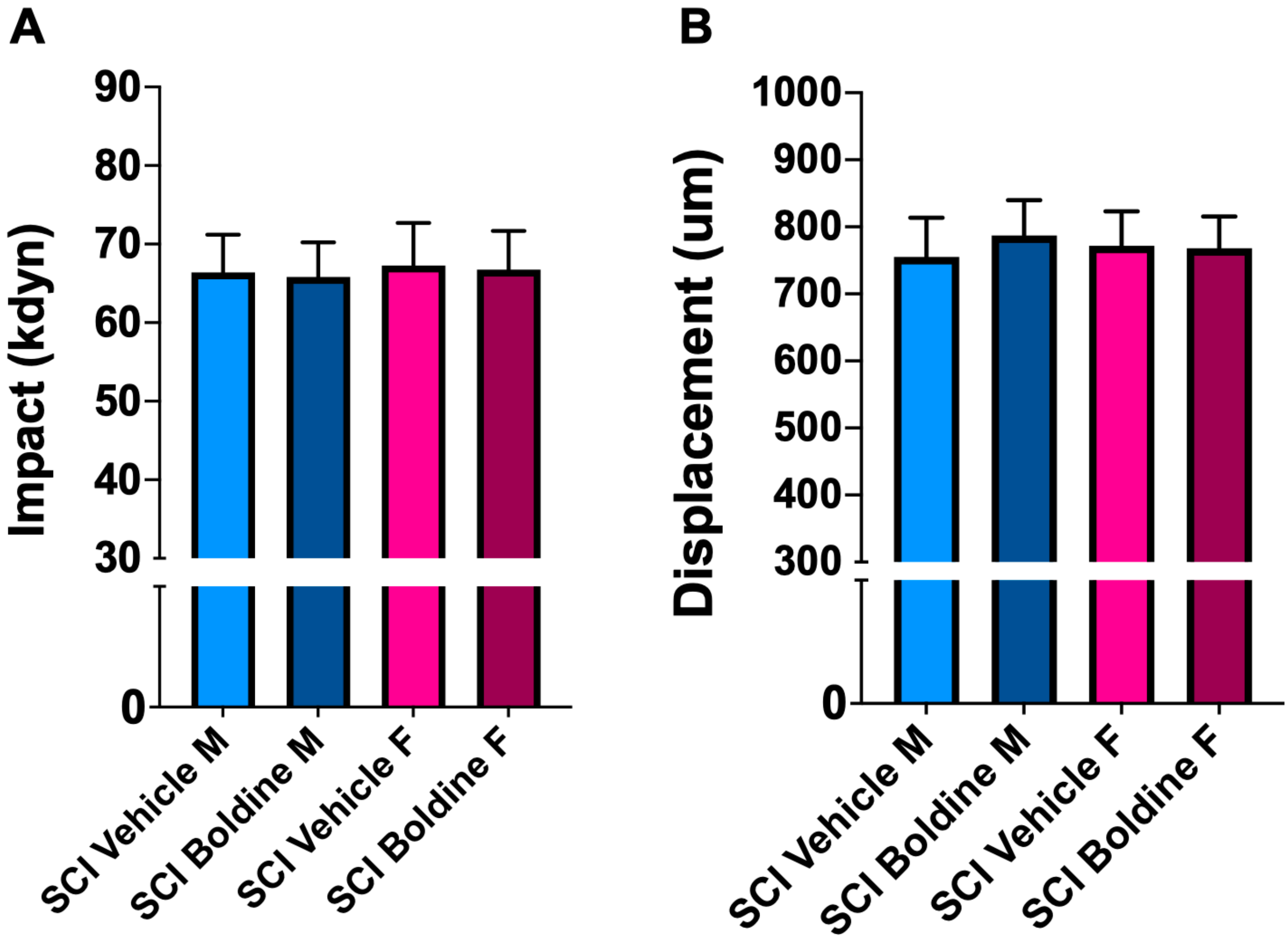


Supplemental Figure 2. Actual impact force (kdyne) and spinal cord displacement during impact (μm) are shown. Data are shown as mean ± SEM.

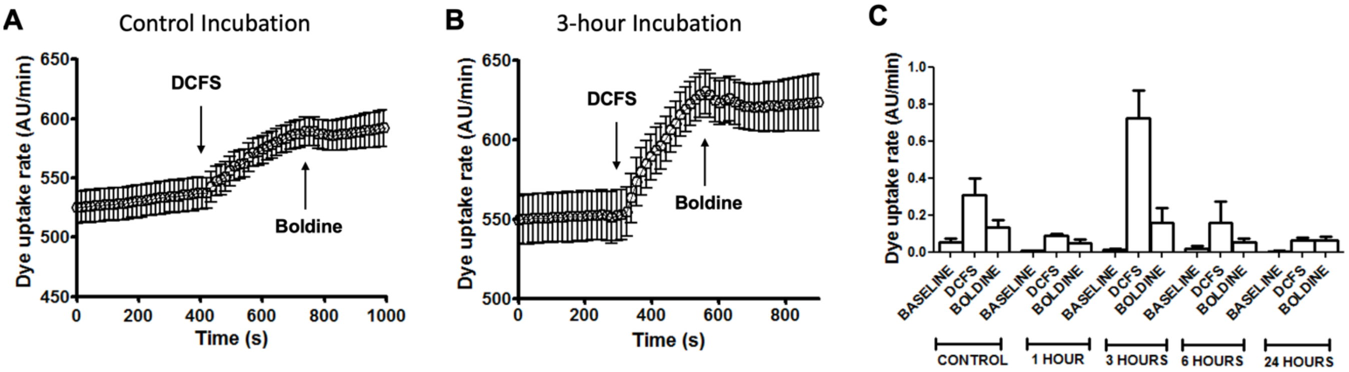


Supplemental Figure 3. Boldine blocks hemichannels expressed by activated spinal cord astrocytes. Spinal cord astrocytes were cultured under control conditions or treated with 10 ng/ml TNF-α plus 10 ng/ml IL-1β 1 at different time periods and dye uptake measured as fluorescence intensity in Arbitrary Units (AU) was evaluated under basal conditions, after exposure to extracellular divalent cation-free solution (DCFS) followed by the application of 50 μM boldine at the times denoted by the arrows in (A) and (B). Curve of fluorescence intensity over time for dye uptake rate was calculated for each condition (C). Each value represents the mean ± SEM of three independent experiments. 30 cells were recorded per experiment.

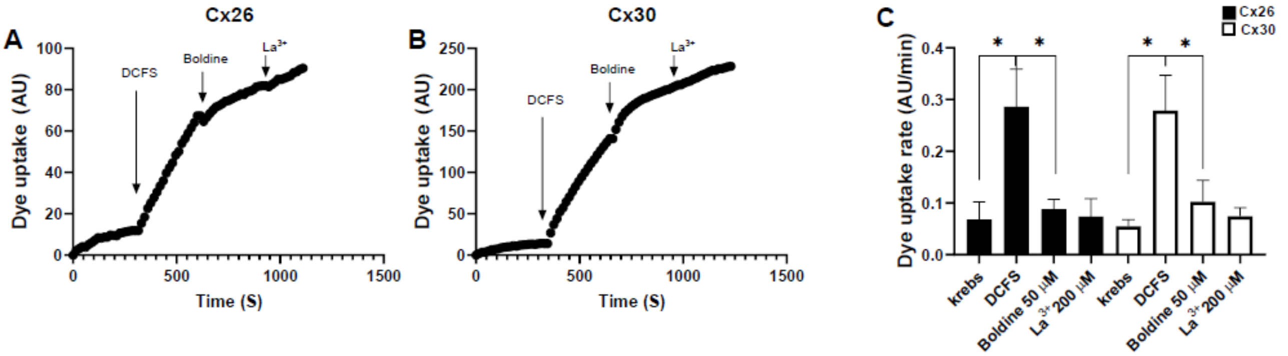


Supplemental Figure 4. Boldine blocks Cx26 and Cx30 hemichannels. A) and B) Cx HC activity was assessed by DAPI uptake measure in time-lapse in Krebs solution, in DCFS to increase the open probability of hemichannels, and in DCFS plus 50 μM Boldine. C) DAPI uptake rate in HeLa Cx26 y HeLa Cx30, the application of DCFS increases the slope of the dye uptake curve and the application Boldine in DCFS drastically reduced the dye uptake. N=4, 30 cells were recorded for each experiment, values are presented as the mean ± SEM. * $p<0.05$. Tukey´s multiple comparisons test.

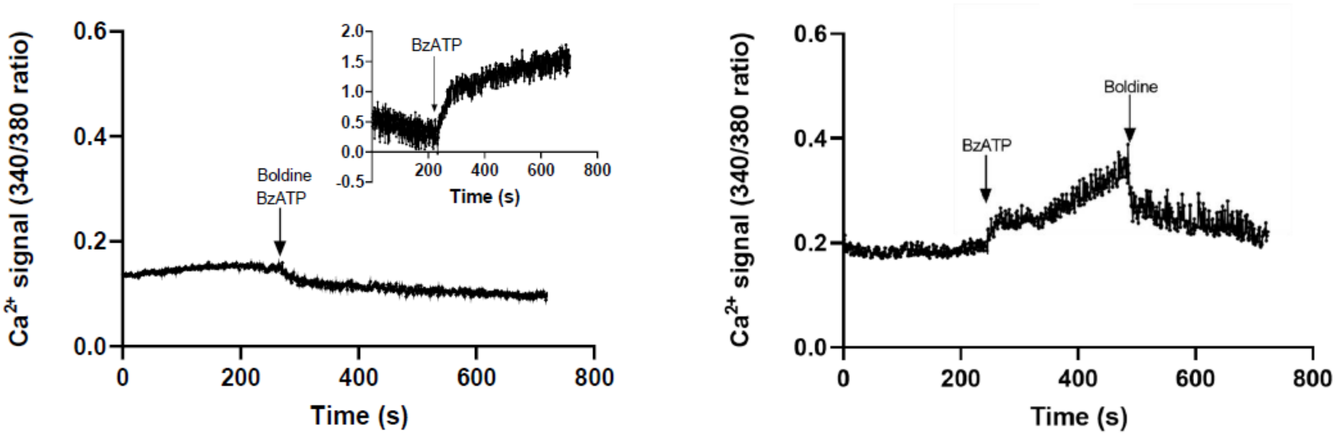


Supplemental Figure 5. Boldine blocks $P2X_7R$. The activity of $P2X_7R$ was evaluated in HeLa cells transiently transfected with mouse $P2X_7R$ -EGFP. Cells were loaded with Fura-2 and intracellular calcium signal was evaluated. Upon treatment with 100 μM benzoyl ATP (inset in the left panel) a calcium signal increase was evident. In some experiments 50 μM boldine (arrows) was added simultaneously with BzATP (left panel) or after cells were treated with BzATP (right panel). Each plotted point corresponds to the mean value of 20 cells of a representative experiment out of four independent experiments.

**A**

| Replicate_name | # Reads |
|---|---|
| Boldine 14dpi below 1 | 29577726 |
| Boldine 14dpi below 2 | 27141109 |
| Boldine 14dpi below 3 | 28985981 |
| Vehicle 14dpi below 1 | 34843051 |
| Vehicle 14dpi below 2 | 32227751 |
| Vehicle 14dpi below 3 | 33166345 |
| Boldine 14dpi above 1 | 27903238 |
| Boldine 14dpi above 2 | 28534783 |
| Boldine 14dpi above 3 | 28433953 |
| Vehicle 14dpi above 1 | 25604140 |
| Vehicle 14dpi above 2 | 28671075 |
| Vehicle 14dpi above 3 | 28485314 |
| Boldine 28dpi below 1 | 30878958 |
| Boldine 28dpi below 2 | 34594307 |
| Boldine 28dpi below 3 | 31361889 |
| Vehicle 28dpi below 1 | 29028843 |
| Vehicle 28dpi below 2 | 31792657 |
| Vehicle 28dpi below 3 | 32531655 |
| Boldine 28dpi above 1 | 34405197 |
| Boldine 28dpi above 2 | 31988594 |
| Boldine 28dpi above 3 | 29760688 |
| Vehicle 28dpi above 1 | 32382925 |
| Vehicle 28dpi above 2 | 31540095 |
| Vehicle 28dpi above 3 | 29937648 |

**B**

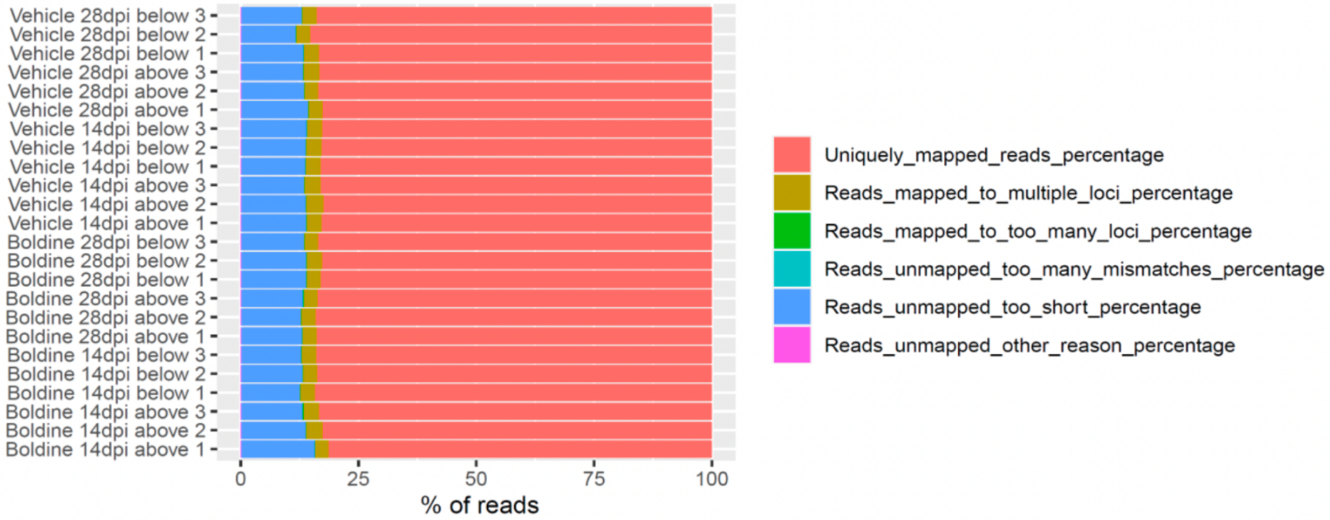


Supplemental Figure 6. Transcriptomic profiling by RNA sequencing. ‘Star’ was used for read alignment to the mouse reference genome. (A) Read counts and (B) alignment efficiencies are shown for each sample.

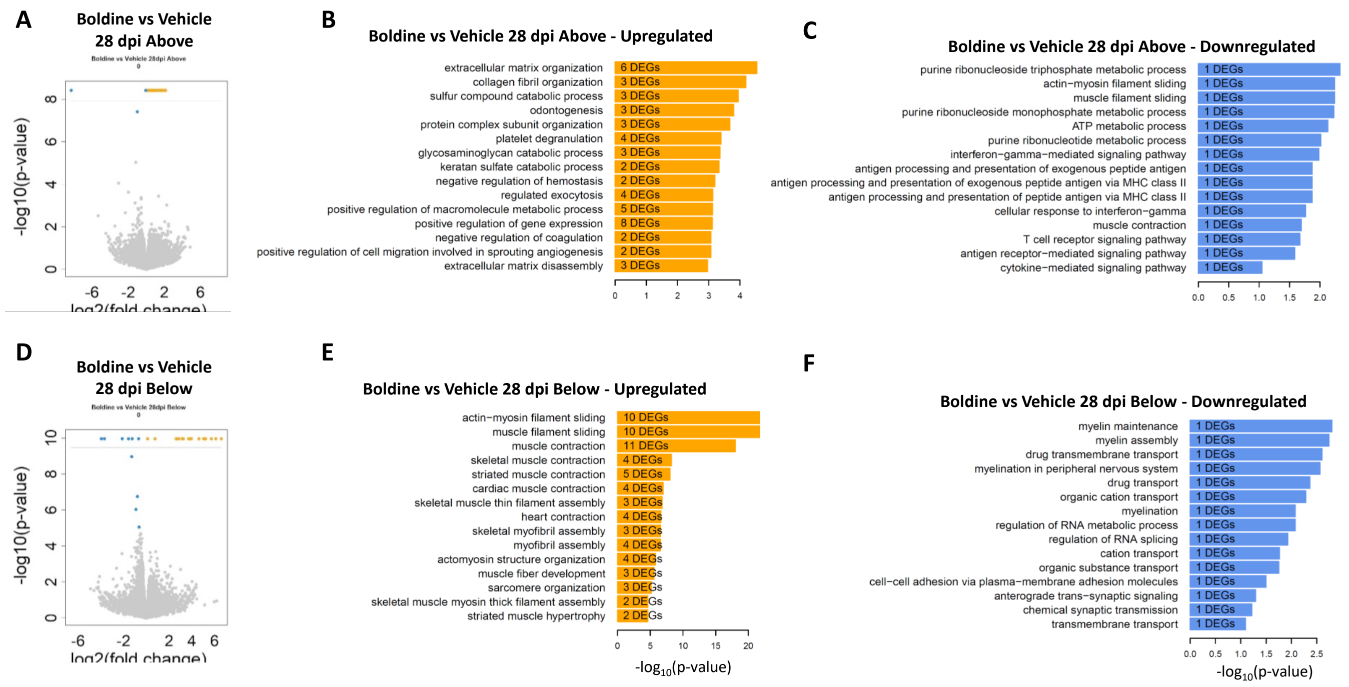


<u>Supplemental Figure 7. Results of analysis by bulk-RNA sequencing of total RNA isolated from spinal cord segments at 28 dpi.</u> Procedures were as described in Methods and the Legend to Figure 5.

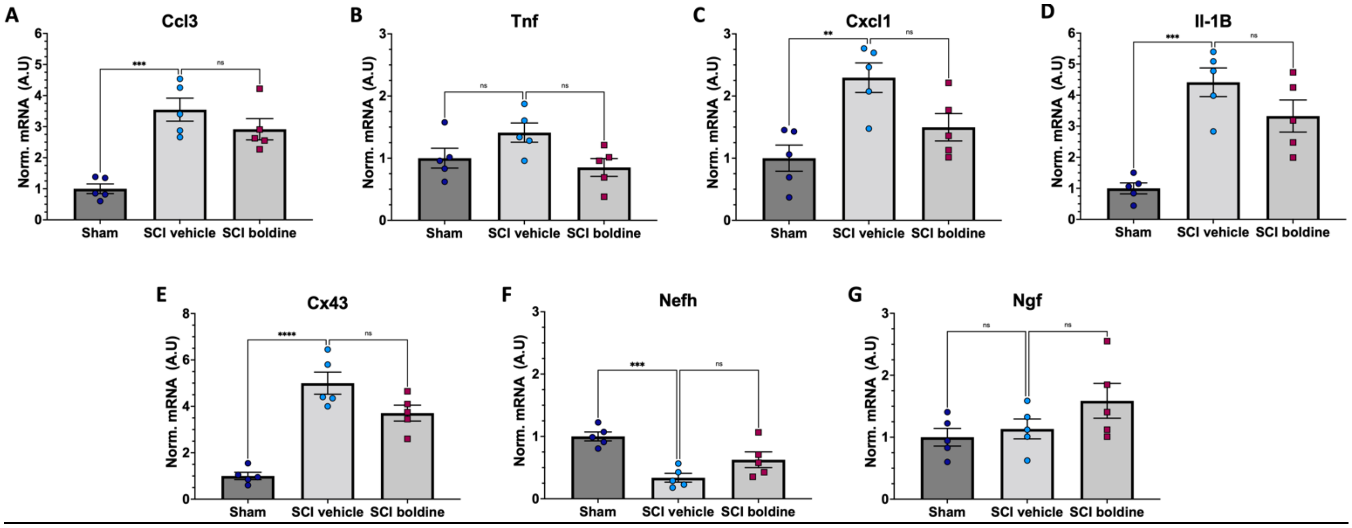


Supplemental Figure 8. Additional RT-qPCR Results. Please refer to Figure 6 and Methods for further details. Data are shown as mean values ± SEM.

See Excel file attached

Supplemental Table 1. Differentially expressed genes between Boldine vs Vehicle above and below at 14 and 28 dpi.

See Excel file attached

Supplemental Table 2. Predicted top 15 Gene Ontology Biological Processes for each list of differentially expressed genes.
